# Slow mitochondria, long life: a phylogenetic comparative analysis in 41 avian species

**DOI:** 10.64898/2026.11.26.769375

**Authors:** Cloe Hadjadji, Suvi Ruuskanen, Julien G.A. Martin, Damien Roussel, Pierre Bize, Antoine Stier

## Abstract

Mitochondrial aerobic metabolism is thought to play a central role in the mechanisms underlying aging. However, because metabolism scales with body mass, which strongly predicts lifespan, disentangling the relationship between mitochondrial metabolism and lifespan from the shared allometric scaling of both traits with body mass remains a persistent challenge. Here, we address this issue by using a phylogenetically controlled, multivariate Bayesian approach to test whether different mitochondrial traits covary with lifespan independently of body mass, across 41 bird species (maximal lifespan: 6–41 years; body mass: 9 g–13 kg). Our results show that, after controlling for body mass, species maximal lifespan was associated with the cellular control of mitochondrial respiration: longer-lived species showed lower respiration under native cellular conditions (i.e. *ROUTINE*), but not at maximum fluxes (i.e. *OXPHOS*). Therefore, longer-lived species used a smaller proportion of their maximal capacity, indicating a larger spare respiratory capacity. Phylogenetic signal was moderate across the four respiration rates, and although mitochondrial traits were tightly correlated within individuals, they showed little evidence of coordinated evolutionary change across species. These results suggest that, in birds, lifespan is linked to how mitochondrial activity is regulated relative to capacity, rather than to maximal performance or efficiency *per se,* and point to the cellular control of mitochondrial respiration as a potential mechanism underlying the evolution of lifespan.

## Introduction

Sustaining life requires a continuous supply of energy, making metabolism a candidate mechanism to explain lifespan variation across species. This reasoning underlies one of the oldest hypotheses in ageing biology, the *rate-of-living theory* (Pearl, 1928), later refined into the *free radical theory of ageing* (Harman, 2003) which predicts that higher mass-specific metabolic rates increase mitochondrial reactive oxygen species (mtROS) production and oxidative damage, and thereby shorten lifespan. Larger species live longer and simultaneously show lower mass-specific metabolic rates than smaller ones (Speakman, 2005). Yet since body mass predicts both traits, this pattern alone does not show that metabolism drives lifespan (Kozłowski, Konarzewski, & Czarnoleski, 2020; Burger, 2024). Moreover, numerous exceptions exist where metabolic rate and lifespan do not covary as predicted, even after accounting for body mass (Speakman, 2005; de Magalhães, Costa, & Church, 2007). Evidence linking oxidative damage to lifespan is also inconsistent (Speakman & Selman, 2011), suggesting that the relevant signal may lie deeper, at the level of the mitochondrion itself. As the site of both ATP synthesis and mtROS production, mitochondria offer a finer scale to test whether the diversity in lifespan and body mass across species is mirrored by variation in mitochondrial aerobic metabolism.

Mitochondria produce most cellular ATP through oxidative phosphorylation (*OXPHOS*). Electrons derived from substrate oxidation flow along the electron transport system (ETS), and the associated energy release drives proton pumping across the inner mitochondrial membrane, generating an electro-chemical gradient (the proton-motive force) used by ATP synthase to produce ATP (Nicholls, 2013). Yet part of the gradient is dissipated as heat when protons leak back into the matrix independently of ATP synthase (*LEAK*), uncoupling substrate oxidation from ATP synthesis, and a fraction of electrons escapes the ETS, mainly at complexes I and III, leading to the formation of mtROS. Mitochondrial coupling efficiency, defined as the extent to which substrate oxidation is coupled to ATP synthesis, depends mainly on the balance between *OXPHOS* and *LEAK*, but also on the electron entry point (i.e. complex I vs. complex II), since complex II does not pump protons (Brand, 2005). Coupling efficiency is likely to be under evolutionary pressure and may underlie variation in organism performance and lifespan (Koch et al., 2021). The *uncoupling to survive hypothesis* stipulates that mild uncoupling may extend lifespan by lowering the proton-motive force and thereby mtROS production (Brand, 2000), and has received mixed empirical support (Speakman et al., 2004; Gates et al., 2007). As *OXPHOS* and *LEAK* share the same proton-motive force, they are functionally linked, and their covariation may reflect common evolutionary history or physiological coupling within species.

Independent of any link to lifespan, mitochondrial traits already show well-established allometric relationships with body mass. Foundational work demonstrated that mitochondrial proton leak scales negatively with body mass across mammals, such that larger species show lower proton conductance than smaller ones (Porter & Brand, 1993). This pattern has been later extended to birds (Brand, Turner, Ocloo, Else, & Hulbert, 2003) and shown to be driven partly by differences in mitochondrial membrane surface area and phospholipid fatty acid composition. Larger species tend to have more monounsaturated membranes that are intrinsically less permeable to protons (Brand et al., 2003). Proton leak and phosphorylating respiration tend to decline with body mass, but as this decline is stronger for *LEAK* than *OXPHOS*, larger species typically also show higher mitochondrial coupling than smaller ones (Boël, Romestaing, Voituron, & Roussel, 2019). Recent work has refined this knowledge, showing that the allometry of coupling efficiency depends on metabolic state (Boël et al., 2019) and tissue type (Barbe, Watson, Roussel, & Voituron, 2023). Mitochondria therefore do not merely scale passively with body size but appear to evolve along their own trajectory, tuned to the specific metabolic demands of the tissue and species in question.

From a life-history or pace-of-life perspective, an analogous set of predictions can be made for lifespan. Short-lived species are expected to favour rapid resource acquisition and immediate energetic output over long-term self-maintenance. Hence, they might show a higher *OXPHOS*, with proportionally lower *LEAK* (i.e., tighter coupling), and greater electron entry through Complex I, which yields more ATP per unit substrate at the cost of higher mtROS production. Current empirical support for a link between mitochondrial traits and species lifespan, independent of body mass, remains limited (Lanza & Nair, 2010). Across 21 bird species, mtROS production and cardiolipin content have been shown to correlate with interspecific variation in lifespan (Delhaye et al., 2016). In marine bivalves, divergences in the control of mitochondrial respiration, rather than in maximal respiratory capacity *per se* were reported to be associated with lifespan variation across 4 species (Rodríguez, Hakkou, Hagen, Lemieux, & Blier, 2021). These studies establish that specific features of mitochondrial traits, beyond simple metabolic rate, covary with lifespan. However, to our knowledge, few studies have tested whether associations between mitochondrial traits and lifespan persist once body mass, together with its established relationships with both variables, is considered (Delhaye et al., 2016).

A further challenge is that mitochondrial traits are usually measured in vitro, on isolated mitochondria or permeabilised tissues or cells, at maximal capacities that are likely rarely attained in vivo, whereas measurements in intact cells provide greater physiological relevance (Divakaruni & Jastroch, 2022). Avian red blood cells possess functional mitochondria (Stier et al., 2013), which allows such measurements from small, minimally invasive blood samples, including in small-bodied and protected species (Stier et al., 2017). Birds thus offer a particularly suitable system to test how mitochondrial traits relate to lifespan (Stier, Massemin, Zahn, Tissier, & Criscuolo, 2015).

In the present study, we investigated how multiple mitochondrial traits vary as a function of species maximum lifespan across 41 avian species spanning a wide range of body masses and lifespans, while explicitly accounting for body mass and phylogeny. Specifically, we predicted that, independent of mass, longer-lived species would show: (i) lower oxidative control efficiency (*OxCE*), achieved through elevated *LEAK*, reduced *OXPHOS* for a given *LEAK*, or a combination of both, consistent with the *uncoupling-to-survive* hypothesis and in contrast to the lower *LEAK* and higher coupling of larger species; and (ii) a lower relative contribution of Complex I to maximal respiration (*OXPHOS_CI_* / *OXPHOS_CI+CII_*), reflecting reduced reliance on the primary site of mtROS production. We predicted the opposite in shorter-lived species, in which greater electron entry through Complex I may be favoured because the resulting gain in ATP-synthesising capacity outweighs the fitness cost of elevated mtROS production over a comparatively shorter lifespan. Because our dataset includes multiple species that share ancestry to varying degrees, we also tested whether each trait carries a phylogenetic signal, i.e. how conserved or evolutionarily labile it is across species, using phylogenetically controlled analysis. For each pair of these traits we also tested (i) whether their covariation reflects a phylogenetic correlation, indicating correlated evolutionary change, through a shared genetic or regulatory basis (e.g., pleiotropy or co-regulation of the underlying genes) or a (ii) non-phylogenetic correlation, for instance through direct mechanistic coupling within the electron transport system. Together, these analyses allowed us to determine whether any lifespan associations are trait-specific and independently evolvable, or instead reflect broader, evolutionarily or mechanistically linked covariation among mitochondrial traits.

## Materials and methods

### Choice of biological material

Avian blood cells have a low mitochondrial metabolism compared to other tissues (Casagrande et al., 2023), but their respiration correlates to mitochondrial respiration in other tissues (pectoral muscle: Stier et al., 2017; brain: (Stier, Monaghan, & Metcalfe, 2022); pectoral muscle and brain: Casagrande et al., 2023), to whole-body metabolic rate (Casagrande et al., 2023) and to organismal performance such as flight speed (Ton, M. Gillings, Pacheco-Fuentes, Stier, & Griffith, 2025; Oefele, Hau, Ruuskanen, & Casagrande, 2026). While these relationships between mitochondrial respirations in avian blood cells with other traits are usually modest in magnitude, and in some cases context-dependent, it demonstrates their relevance up to the organismal scale and for a comparative study (Delhaye et al., 2016, current study).

### Animal sampling

We gathered blood samples from 219 individuals from 41 species between 2018 and 2025 (ESM, Table S1). All birds were adults being mostly caught around the reproductive season, with in most cases no information on the exact breeding status, sex and age. While mitochondrial aerobic metabolism in avian blood cells has been shown to vary with reproductive (Stier, Bize, Hsu, & Ruuskanen, 2019), sex (Stier, Monaghan, & Metcalfe, 2022; Cossin-Sevrin et al., 2025; Beccardi et al., 2026) and age (Stier, Monaghan, & Metcalfe, 2022; Cossin-Sevrin et al., 2025; Beccardi et al., 2026), and body mass and lifespan can also vary between sexes for a given species, these inter-individual variations are expected to be relatively minor compared to inter-species variation. We sampled two captive species (Japanese quail and chicken; authorization APAFIS#54114-2025030516541037 v4) and 39 wild species. Wild animals were sampled as part of regular mistnetting sessions and population monitoring in Turku, Finland (32 species; authorization (ESAVI/5718 and VARELY/924), the French Southern Territories (6 species; authorizations APAFIS#16465-2018080111195526 v4 and Arrêté TAAF A-2018-118) and Switzerland (1 species, authorization 26552). Blood samples were collected from the brachial vein using heparinized capillaries or syringes for all bird species except Alpine swifts where blood samples were collected from the foot vein using heparinized microvette tubes (Sarstedt, Germany). All samples were kept on ice until laboratory analysis within 24 hours maximum (Stier et al., 2017).

### Mitochondrial aerobic metabolism measurements

All the measurements of mitochondrial aerobic metabolism were performed using an O2k high-resolution respirometer (Oroboros Instruments, Innsbruck, Austria). Measurements were performed on blood cells following the protocol of Stier et al. (2019), at 40°C, except for king penguin that were measured at 38°C according to their normal body temperature and previous studies on mitochondrial metabolism (Stier et al., 2017). Between 20 and 50 µL of whole-blood was suspended in Mir05 buffer (EGTA [0.5 mmol l^−1^], MgCl2 [3 mmol l^−1^] , K-lactobionate [60 mmol l^−1^] , taurine [20 mmol l^−1^] , KH2PO4 [10 mmol l^−1^] , HEPES [20 mmol l^−1^] , sucrose [110 mmol l^−1^] , free fatty acid bovine serum albumin [1 g.L-1], pH 7.1; Sigma-Aldrich, St. Louis, MO, USA), washed once by centrifugation for 2 minutes at 1000g, and resuspended in pre-warmed Mir05 in the O2k chamber. In total we measured sequentially five distinct respiration rates including: (1) *ROUTINE*: endogenous cellular respiration rate of intact cells before permeabilization with digitonin [10 µg mL^−1^]; (2) *OXPHOS_CI_*: the maximum phosphorylating respiration rate fuelled with exogenous substrates of complex I (i.e., malate [2 mmol l^−1^], pyruvate [5 mmol l^−1^]) and ADP [1.25 mmol l^−1^]; (3) *OXPHOS_CI+CII_*: the maximum phosphorylating respiration rate fuelled with exogenous substrates of complex I and II (i.e., adding succinate [5 mmol l^−1^]); (4) *LEAK* respiration when ATP synthesis is inhibited with 2.5 mmol l^−1^ of oligomycin, and therefore respiration is mostly driven by mitochondrial proton leak; (5) *ROX*: non-mitochondrial respiration after complex III inhibition with 5 mmol l^−1^ of antimycin A. *ROX* was then subtracted from other respiration rates to remove the influence of non-mitochondrial respiration. Respiration rates were expressed relative to cell number (i.e. pmol O_2_/s/10^6^ cells), which was evaluated with an automatic cell counter (Bio-Rad TC20 or Curiosis FACSCOPE B). From these respirations rates we calculated three mitochondrial flux control ratios, specifically: *ROUTINE/OXPHOS_CI+CII_* indicating the proportion of maximal capacity being used under endogenous cellular conditions, *OXPHOS_CI_*/*OXPHOS_CI+CII_* indicating the relative contribution of complex I to overall *OXPHOS* respiration, and the OXPHOS control efficiency *OxCE* ((*OXPHOS_CI+CII_* - *LEAK*)/*OXPHOS_CI+CII_*) indicating the proportion of mitochondrial respiration being linked to ATP synthesis. Unfortunately, the more direct indicator of mitochondrial efficiency (i.e. ATP/O ratio) cannot be reliably estimated with the available methods because of the low respiration rates in avian blood cells.

### Lifespan and body mass records

Maximum lifespan and average species body mass were extracted from AnAge database for 39 species (De Magalhães & Costa, 2009). There was no information in this database for Lesser sheathbill (*Chionis minor*) and White-chinned petrel (*Procellaria aequinoctialis*), but thanks to the long-term ringing and monitoring conducted by the French Polar Project #109, we obtained very robust estimates for both species (Barbraud et al., 2011). Maximum lifespan and average body mass used in this study are available in ESM Table S1.

### Comparative analysis

All statistical analyses were performed using R version 4.6.1 (R Core Team, 2026). To examine the relationship between mitochondrial respiration rates and lifespan across species, we used Bayesian phylogenetic mixed models implemented in the MCMCglmm package (Hadfield, 2010). All mitochondrial traits were scaled prior to analysis to facilitate comparison of estimates. For each trait, we modelled mitochondrial respiration as a function of log-transformed body mass and residual lifespan. We obtained residual lifespan by extracting residuals of a log-log regression of maximal lifespan against body mass (*R^2^* = 0.40, F_1,39_ = 26.45, p-value < 0.001), thereby capturing variation in lifespan independent of body mass. This approach is standard in comparative biology and physiology (e.g., de Magalhães et al., 2007; Minias & Podlaszczuk, 2017; Liu et al., 2023). We also performed the present analysis using log-transformed lifespan and log-transformed body mass, which yielded the same results for the effect of lifespan, and present them in the Electronic Supplementary Material (Table S2).

To account for the non-independence of species data due to shared evolutionary history, we included a random effect of species identity linked to a phylogenetic tree. The phylogenetic tree used in this study was the Aves 1.6 tree (McTavish et al., 2025). Since Aves 1.6 provides a single consensus tree, no topology averaging was required. The tree was pruned to our 41 study species using the *keep.tip* function from the ape package (ESM Figure S1, Paradis et al., 2024). Following pruning, ultrametricity was enforced using the *chronopl* function (lambda = 0) to correct for minor numerical errors introduced during pruning. The phylogenetic relatedness matrix was then scaled and inverted using the *inverseA* function from the MCMCglmm package and included directly in the models as a *ginverse* term linking the species identity to their phylogeny (Hadfield, 2010). Since we have repeated measurements per species, we also included an additional species identity random effect but not link to the phylogeny to capture residual among-species variance not attributable to phylogeny, allowing the model to partition variance into phylogenetic and non-phylogenetic components.

Our four mitochondrial traits (*ROUTINE, OXPHOS_CI_, OXPHOS_CI+CII_,* and *LEAK*) have strong biological and functional interdependence, and thus they were modelled jointly in a single multivariate MCMCglmm model. It allowed us to estimate trait-specific fixed effects while accounting for phylogenetic and residual covariance among traits. This model included a trait-specific intercept and trait-specific slopes for log body mass and residual lifespan, an unstructured phylogenetic random-effect structure by trait, an unstructured species-identity random-effect structure by trait, and an unstructured residual (co)variance structure by trait, allowing correlations among traits to be estimated at each variance level. Because the remaining mitochondrial traits are compound ratios (*ROUTINE*/ *OXPHOS_CI+CII_*, *OXPHOS_CI_*/*OXPHOS_CI+CII_* and *OxCE*), they cannot be directly comparable in a joint covariance framework. Therefore, they were modelled separately using univariate models with the same phylogenetic and species-identity random-effect structure described above.

For the univariate models, we specified parameter-expanded priors for the random effects (V = 1, ν = 1, α.µ = 0, α.V = 1000), which place diffuse, weakly informative priors on the variance components and improve mixing when sample sizes are small; a non-informative prior was used for the residual variance (V = 1, ν = 0.002). For the multivariate model, we specified standard inverse-Wishart priors for the unstructured (co)variance matrices at each random-effect level (V = diag(4) × 0.5, ν = 4), reflecting the number of jointly-modelled traits. We additionally ran models under alternative prior specifications (varying the informativeness of ν and V), posterior point estimates and credible intervals were consistent across specifications, mixing and effective sample size were best under the priors reported here. Univariate models were run for 2,500,000 iterations with a burnin of 500,000 and a thinning interval of 1,000, resulting in 2,000 posterior samples. The multivariate model was run for 1,500,000 iterations with a burnin of 200,000 and a thinning interval of 500, resulting in 2,600 posterior samples. Convergence was assessed by visual inspection of trace plots, examination of autocorrelation functions, and Heidelberger-Welch stationarity and half-width tests.

For each trait, we additionally calculated phylogenetic signal (λ) as the proportion of total variance attributable to the phylogenetic random effect relative to the sum of the phylogenetic, species-identity, and residual variance components (De Villemereuil, 2019), reporting the posterior median and 95% HPDI. To test whether mitochondrial traits evolve together independently of their shared relationships with body mass and residual lifespan, we extracted the posterior phylogenetic, species-identity (non-phylogenetic), and residual (within-species, individual levels) (co)variance matrices from the unstructured random-effect terms of the multivariate model and converted these to correlation matrices. We report the posterior median and 95% HPDI for each pairwise trait correlation at three levels: the phylogenetic level (reflecting evolved covariance among traits attributable to shared ancestry, independent of the fixed effects), the species level (reflecting repeatable among-species covariation not explained by phylogenetic relatedness), and the residual level (reflecting within-species covariation among individuals). We additionally examined the joint posterior distributions of the phylogenetic and species-identity variance components to assess the degree of statistical trade-off between these two random effects, which share the same grouping factor (species identity).

We report posterior mean and 95% highest posterior density intervals (HPDI) for each fixed-effect parameter, and consider strong evidence for an effect when the 95% HPDI do not overlap zero and when *P_MCMC_*< 0.05. Since variance parameters and variance ratio (i.e., λ) are positive by definition, we considered HDPI including values below 0.001 to provide weak evidence of an effect. A correlation was considered to have strong evidence when its 95% HPDI excluded zero.

## Results

The relationships between mitochondrial traits and avian body mass are shown in Figure 1, measurements were performed on 41 bird species (219 individuals), spanning 21 families and 9 orders (Anseriformes, Apodiformes, Charadriiformes, Galliformes, Passeriformes, Piciformes, Procellariiformes, Sphenisciformes, Strigiformes), and ranging from 9 g to 13 kg (see full details in Table S1).

**Figure 1.**
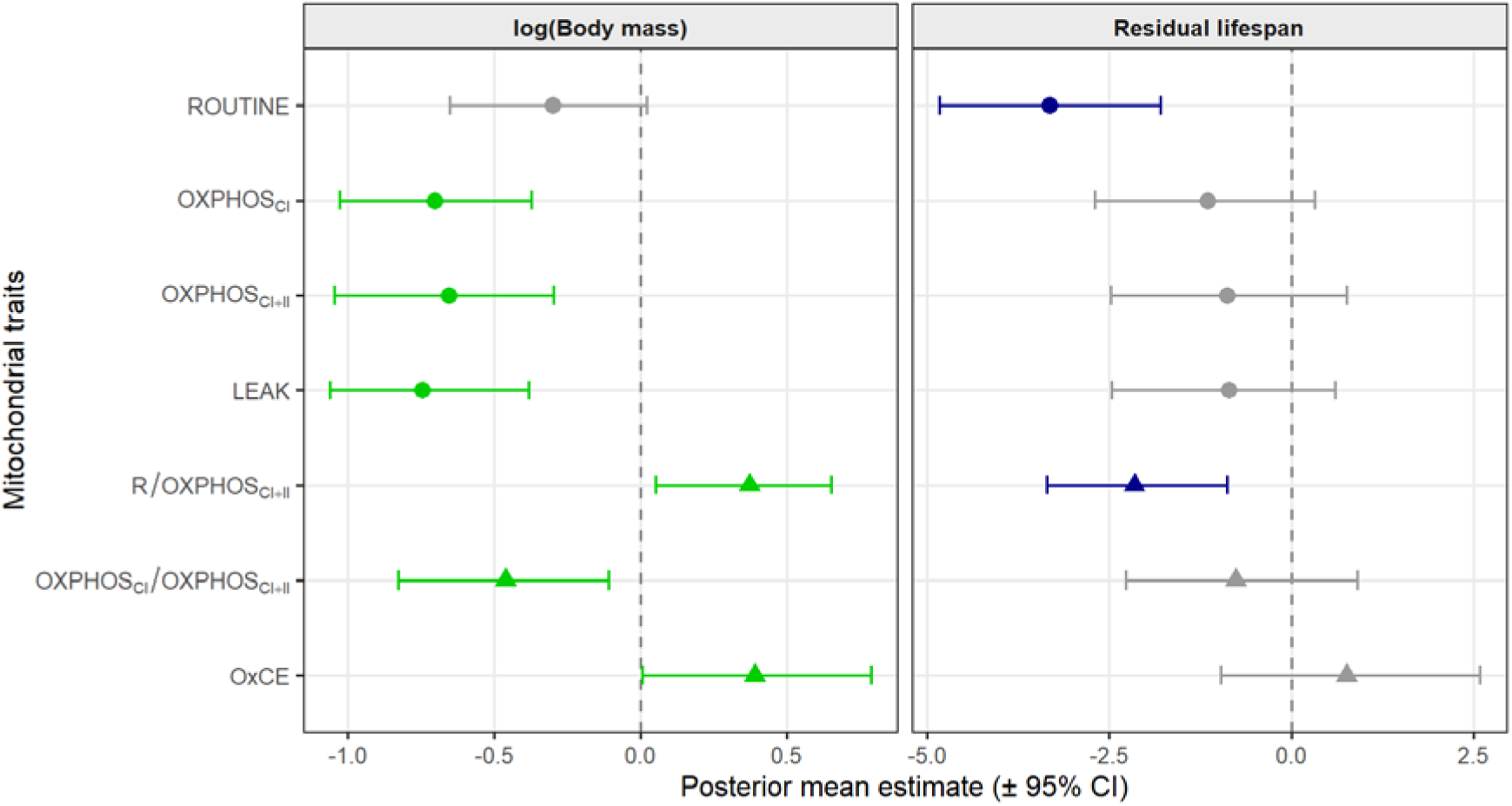
Relationships between mitochondrial traits and species: (A) body mass, and (B) residual maximum lifespan. Points show posterior mean estimates and horizontal bars their 95% credible intervals (CI) from phylogenetic MCMCglmm models (Aves 1.6 tree). Circles denote estimates from the multivariate model for respiration rates (ROUTINE, OXPHOSCI, OXPHOSCI+II, LEAK) modelled jointly; triangles denote estimates from univariate models fitted separately for each mitochondrial flux control ratio (R/OXPHOSCI+II, OXPHOSCI/OXPHOSCI+II, OxCE). Effects with strong evidence (95% HPDI excludes zero) are shown in green (body mass) or dark blue (residual lifespan); effects with weak or no evidence (95% HPDI overlaps zero) are shown in grey.

After accounting for phylogenetic relatedness, the relationship with body mass showed negative allometry for *ROUTINE* (marginal)*, OXPHOS_CI_, OXPHOS_CI+CII_, LEAK* and *OXPHOS_CI_/OXPHOS_CI+CII_*(Figure 1, ESM Table S3), indicating that larger species have lower mitochondrial respiration rate and lower reliance of complex I. In contrast, *R*/*OXPHOS_CI+CII_* sand *OxCE* showed evidence for positive allometry (Figure 1, ESM Table S3), indicating that larger species use a greater share of their maximal mitochondrial capacity under endogenous cellular conditions and presented higher mitochondrial control efficiency.

Analyses of the relationships of mass-corrected residual lifespan with mitochondrial traits revealed a clear negative association for *ROUTINE* and *R/OXPHOS_CI+CII_* only (Figure 1 and 2). Longer-lived species, relative to their mass, show lower *ROUTINE* respiration and use a smaller share of their maximal mitochondrial capacity under endogenous cellular conditions. All other mitochondrial traits were not significantly associated with residual lifespan after controlling for direct and indirect effect of body mass.

**Figure 2.**
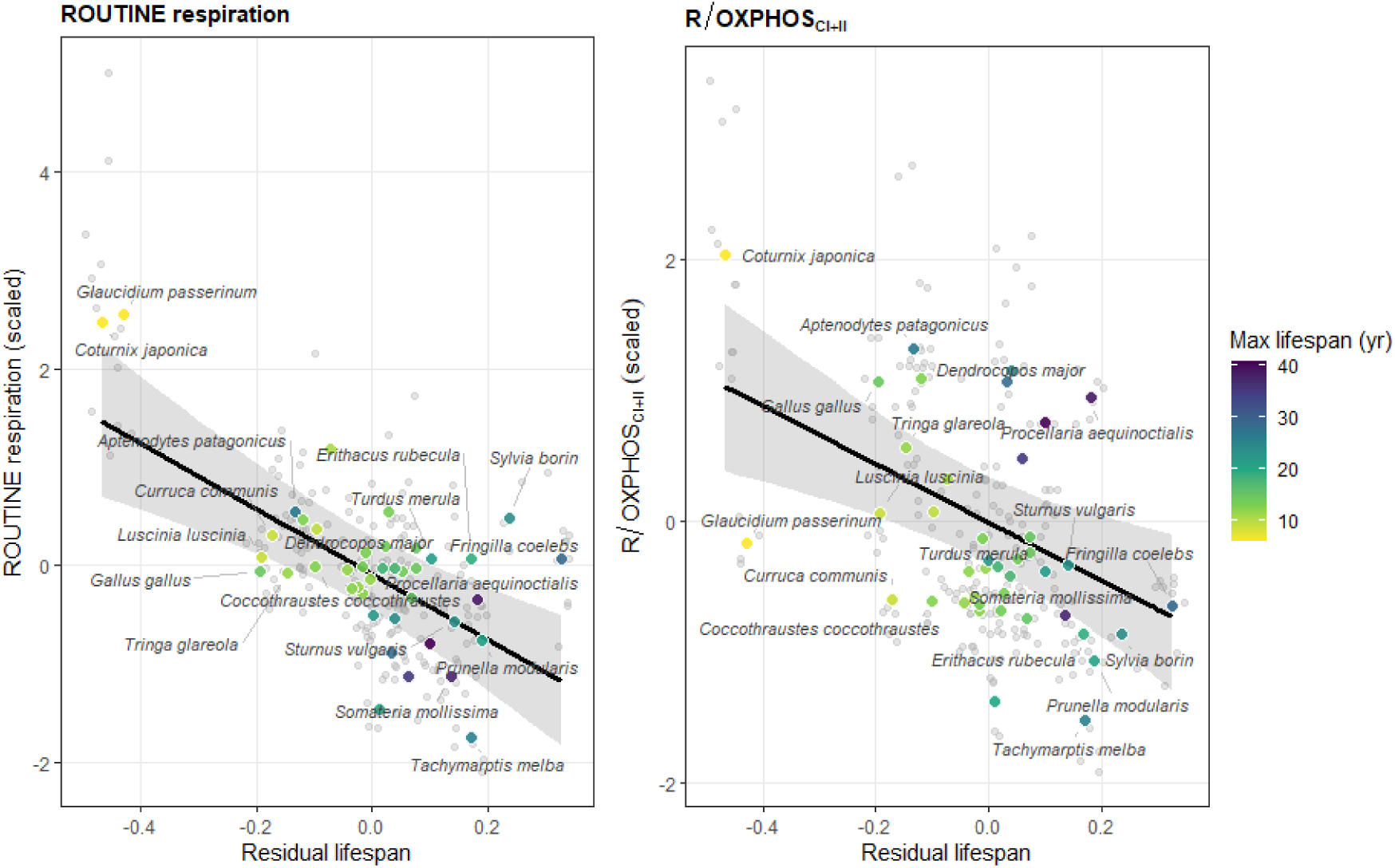
Relationship between residual lifespan and (A) mitochondrial respiration under endogenous cellular conditions, (B) the flux control ratio estimating the proportion of maximal mitochondrial capacity being used under endogenous cellular conditions. Coloured points show species averages, coloured from yellow to dark blue according to maximum lifespan (6–41 years); grey points show individual measurements included in the underlying MCMCglmm models (multivariate model for ROUTINE, univariate model for R/OXPHOSCI+II ratio). The black line and shaded ribbon show the posterior prediction and 95% credible interval at mean body mass. Relationships for the remaining, non-significant traits are shown in the supplementary material.

Phylogenetic signal was low to moderate across mitochondrial traits (Figure S2). For the four mitochondrial respiration rates modelled jointly (*ROUTINE, OXPHOS_CI_, OXPHOS_CI+CII_, LEAK*), estimates were statistically supported (posterior median λ = 0.32–0.37, all HDPI > 0.001), indicating a significant phylogenetic signal. In contrast, phylogenetic signal estimated univariately for the mitochondrial flux control ratios (*OxCE, R/OXPHOS_CI+CII_, OXPHOS_CI_/OXPHOS_CI+CII_*) was lower, with all 95% HPDI close to 0 (posterior median λ = 0.06–0.27), indicating little to no evidence of phylogenetic signal for these traits.

We then examined how the four respiration rates (*ROUTINE*, *OXPHOS_CI_*, *OXPHOS_CI+II_*and *LEAK*) covary, by partitioning their pairwise correlations into a phylogenetic component (covariation attributable to shared ancestry), a non-phylogenetic species-level component (covariation among species not explained by relatedness), and a residual component (covariation among individuals within species; Figure S3A). Phylogenetic correlations among *ROUTINE, OXPHOS_CI_, OXPHOS_CI+II_,* and *LEAK* were generally weak, and only *OXPHOS_CI_* and *OXPHOS_CI+II_* showed a statistically supported phylogenetic correlation (*cor_phylo_* = 0.58 [0.17; 0.84]). Non-phylogenetic species-level correlations followed a closely matching pattern: *OXPHOS_CI_* and *OXPHOS_CI+II_* again showed the only evident correlation among species not attributable to phylogenetic relatedness (*cor_species_* = 0.54 [0.14; 0.81]), while all other pairwise correlations remained weak (*cor_species_* = 0.13-0.36). In contrast, within-species (residual) correlations among the same four traits all had strong evidence (HPDI not overlapping zero) and were large (*cor_residuals_*= 0.45-0.88; ESM Figure S3A). For all four mitochondrial traits, total phenotypic variance was partitioned approximately equally among phylogenetic (posterior modes: 31– 36%), non-phylogenetic species (27–31%), and within-species residual (31–40%) components, although HPD intervals were wide and broadly overlapping (Figure S3B), reflecting limited power to separate phylogenetic from non-phylogenetic species variance with 41 species.

## Discussion

This study investigates the links between 7 mitochondrial traits and maximal lifespan using a phylogenetically controlled comparative analysis in 41 bird species. Results show that maximal lifespan was associated with the cellular control of mitochondrial respiration (*R/OXPHOS_CI+CII_*), rather than with maximal respiratory capacity (*OXPHOS_CI+II_*), proton leak (*LEAK*), or coupling efficiency estimate. Phylogeny also accounted for a significant part of the variation in mitochondrial respiration rates, but not in the flux control ratios, suggesting that absolute respiratory capacity is evolutionarily more conserved than the relative balance of mitochondrial traits, a distinction we return to below.

The negative association between *ROUTINE* respiration and lifespan indicates that long-lived species maintain lower endogenous mitochondrial activity, independent of body mass. They also use proportionally less of their maximal capacity at rest (lower *R/OXPHOS_CI+CII_*), indicating a larger reserve respiratory capacity (also called spare respiratory capacity), compared to short-lived species (Desler et al., 2012). Reserve respiratory capacity represents the difference between resting and maximal respiratory output and provides a potential buffer when ATP demand increases. Depletion of this reserve has been linked to cellular stress vulnerability and age-related dysfunction in mammals, including in neurons and cardiomyocytes (Desler et al., 2012; Marchetti, Fovez, Germain, Khamari, & Kluza, 2020). A larger reserve might therefore constitute a protective and adaptive trait that scales with the evolution of longer lifespan across species. This raises the possibility that longer-lived species are potentially better equipped to adapt quickly when facing stressful conditions that require a rapid rise in ATP demand. The lower *ROUTINE* respiration of long-lived species further suggest that these species may generate less mtROS under basal physiological conditions, assuming that mtROS production scales positively with *in vivo* basal electron flux (Barja, 2013). However, because we did not measure mtROS production, nor demonstrated a positive link between mtROS production and *ROUTINE* respiration, this interpretation remains to be formally tested. This interpretation is nonetheless consistent with prior work in using avian red blood cells showing that long-lived species produce less mtROS than short-lived ones (Delhaye et al., 2016). Altogether, our results offer two complementary, non-mutually exclusive explanations (preserved bioenergetic reserve and reduced ROS generation in long-lived species) by which lower endogenous mitochondrial activity could plausibly be linked to extended lifespan. Our results also converge to some extent with a comparative study in four marine bivalve species, where the longest-lived species showed reduced control by electron-entry pathways (Complex I, Complex II) and increased control by Complex IV (Rodríguez et al., 2021), pointing out that how respiration is regulated along the electron-transport system may be more relevant to lifespan than maximal respiratory capacity alone.

While we predicted that long-lived species may exhibit lower *OXPHOS* control efficiency (*OxCE*), because mildly uncoupled mitochondria generate less mtROS (i.e., *“uncoupling-to-survive“* hypothesis; Brand, 2000), we found no such evidence. We also found no evidence for short-lived species to maximize ATP efficiency through a preferential use of complex I for electron entry. These findings should nevertheless be interpreted cautiously because *OxCE* and *OXPHOS_CI_/OXPHOS_CI+CII_* are indirect measures related to mitochondrial efficiency, but do not directly quantify ATP production per unit of oxygen consumed (ATP/O). Together, it suggests that mitochondrial regulation, rather than capacity or efficiency, may be a recurrent axis linking mitochondrial traits to lifespan across distant lineages. The mechanisms underlying this regulation remain to be determined: lower *ROUTINE* respiration relative to capacity may reflect differences in cellular energy demand, substrate supply, or intrinsic differences in how mitochondrial respiration is regulated, which our design cannot distinguish. Our results also highlight the importance of studying mitochondrial respiration under physiologically relevant conditions, such as here with *ROUTINE* being measured in intact cells with their own substrates and cellular control. Indeed, intact-cell respiration reflects integrated cellular metabolism, whereas permeabilized-cell/isolated-mitochondrial assays provide much greater experimental control but sacrifice physiological context (Divakaruni & Jastroch, 2022).

As expected, we found negative allometric results between body mass and mitochondrial respiration rates, although borderline non-significant for *ROUTINE*. These results agree with previous findings using isolated mitochondria in skeletal muscles of mammals (Boël et al., 2019) and birds (Barbe et al., 2023). It suggests mitochondrial respiration in avian blood cells follow the same pattern as in other tissues. The fact that *OxCE* (average values in red blood cells: 0.84) only marginally scaled with species body mass may be in accordance with results of Barbe et al., (2023) who did not find scaling between Respiratory Control Ratio (RCR = 1/(1-*OxCE*); OxCE_Heart/Muscle_ = 0.78-0.79) and body mass. Yet, *OxCE* showed a marginal positive scaling with body mass, which may be in line with results from Brand et al., (2003) showing that RCR increases with species body mass. To the best of our knowledge, no study so far tested the scaling with body mass of complex I relative contribution (*OXPHOS_CI_*/*OXPHOS_CI+CII_*) nor the cellular control of respiration (*R/OXPHOS_CI+CII_*). Here, we demonstrate that larger species have a lower relative contribution of complex I to maximum OXPHOS capacity. This is consistent with recent findings in mammals showing that NAD-dependent enzymes decline with increasing body mass (Boël, Voituron, & Roussel, 2026), which suggest a reduced contribution of complex I in larger species, as NADH-derived electrons enter the ETC principally through complex I.

The four mitochondrial respiration rates (*ROUTINE*, *OXPHOS_CI_*, *OXPHOS_CI+CII_*, *LEAK*) showed statistically supported moderate phylogenetic signal (λ = 0.32–0.37; Figure S2). This is within but overall lower than the range reported by Barbe et al., (2023) for other mitochondrial traits (approximate range: *h*^2^ 0.25–0.90) but nonetheless indicating that absolute respiration rates carry a detectable, if modest, evolutionary basis across the avian phylogeny. By contrast, the three flux control ratios (*OxCE*, *R*/*OXPHOS_CI+CII_*, *OXPHOS_CI_*/*OXPHOS_CI+CII_*) showed very weak evidence of phylogenetic signal, with moderate point estimates (λ = 0.06–0.27) and all 95% HPD intervals overlapping zero. One of the plausible mechanistic explanation specific to our traits is that because absolute respiration rates scale with mitochondrial density, any phylogenetic signal detected in *ROUTINE*, *OXPHOS_CI_*, *OXPHOS_CI+CII_*, and *LEAK* could partly reflect conserved evolutionary differences in mitochondrial density itself, rather than in the functional properties of the electron transport system *per se.* Flux control ratios, being calculated as the ratio of two rates measured on the same mitochondria, are by construction independent of density and instead capture the relative, qualitative balance of ETS function. The absence of phylogenetic signal in these ratios therefore suggests that their variations may be shaped more by current physiological demands or plasticity than by conserved ancestry.

A similar conclusion can be drawn from the overall weak and largely unsupported phylogenetic correlations among the four respiration rates. These traits appear to evolve largely independently of one another across species rather than showing coordinated co-evolution. Notably, *ROUTINE* was strongly correlated at the residual level with *OXPHOS_CI_*, *OXPHOS_CI+CII_*, and *LEAK* at the individual level, indicating that individuals with higher mitochondrial capacity also tend to operate at a proportionally higher endogenous activity. However, only *ROUTINE*, and the fraction of capacity it represents (*R*/*OXPHOS_CI+CII_*), showed a species-level association with lifespan. This pattern suggests that the physiological coupling observed within species does not necessarily translate into coordinated evolutionary change across species. Species may therefore have diverged in their basal mitochondrial state relative to capacity, partly independently of divergence in maximal capacity itself. Since *ROUTINE* was the only trait strongly associated with lifespan, despite this residual correlation with *OXPHOS_CI_*, *OXPHOS_CI+CII_*, and *LEAK*, the association with lifespan is best interpreted as trait-specific rather than reflecting a broader, coordinated shift in mitochondrial traits. Overall, this means that mitochondrial traits are tightly coupled within species, but largely uncoupled evolutionarily, across species.

The 41 bird species included in this study span a wide range of lifespans, body masses, environments, and ecologies, and represent a substantial phylogenetically controlled comparative dataset of mitochondrial traits. Species at opposite ends of the *ROUTINE*–lifespan relationship tend to differ in broader ecological or life-history strategy. Several of the longest-lived, lowest-activity species in the present dataset are seabirds (i.e., *Procellaria aequinoctialis:* White-chinned petrel, *Pygoscelis papua*: Gentoo penguins), a group classically associated with delayed maturity and low reproductive rate, as well as other long-lived species such as the Alpine swift (*Tachymarptis melba*) or the common eider (*Somateria mollissima*). The shortest-lived, highest-activity species, by contrast, are also among the species with the smallest body size, such as the pygmy owl (*Glaucidium passerinum*) and the Japanese quail (*Coturnix japonica)*. Yet since we use residual lifespan, the position of long versus short-lived species along a slow-fast pace of life continuum cannot be attributed to body mass, suggesting that additional ecological or life-history factors beyond size contribute to their comparatively slow versus fast-paced mitochondrial activity. This pattern is broadly consistent with a wider pace-of-life continuum (Brown, Burger, Hou, & Hall, 2022), in which ecological pressures shaping reproduction and survival may act on the regulation of mitochondrial activity.

In conclusion, we found that maximal lifespan in birds is linked to how mitochondrial activity is regulated relative to capacity, rather than to maximal respiratory capacity or efficiency themselves. This regulatory balance, captured specifically by *ROUTINE* respiration, constitutes a candidate physiological link between mitochondrial traits and lifespan, independent of body mass, and connects to prior evidence linking lower mtROS production to lifespan in birds. Cellular regulatory control may be the more evolutionarily relevant target here. Indeed, *ROUTINE* reflects mitochondrial activity under endogenous substrate supply and normal cellular regulation, unlike measures of electron transport system capacity alone. This underscores the value of intact-cell respirometry for evolutionary and ecological questions. Finally, partitioning trait covariation into phylogenetic, species-level, and residual components revealed that mitochondrial traits are tightly integrated physiologically within species yet evolve largely independently of one another across species. This distinction is rarely resolved in comparative datasets, and one that clarifies why the lifespan signal identified here is trait-specific rather than reflecting a broader, coordinated shift in mitochondrial traits.

## Acknowledgments

We are grateful to the bird ringers from Turku botanical garden, to T. Laaksonen, C. Morosinotto and C. Arzel for providing blood from their study systems in Finland, to the French Polar projects #109, #119 and #1151 for their help in collecting blood from avian species in Crozet Archipelago, and to people helping collecting some of the mitochondrial data (N. Garcin, N. Cossin-Sevrin and C. Bocquet).

## Funding acquisitions

This research was funded by the Turku Collegium for Science and Medicine and a Marie Sklodowska-Curie Postdoctoral Fellowship (#894963) to AS, the Research Council of Finland (grant #293943) to SR and the Swiss National Science Foundation (grant #212971) to PB.

## Authors contributions

**C. Hadjadji:** Writing – original draft, Visualization, Formal analysis, Data curation; **S. Ruuskannen:** Conceptualization, Data collection, Writing – review & editing; **J. G.A. Martin:** Validation, Writing – review & editing; **D. Roussel:** Writing – review & editing; **P. Bize:** Validation, Writing – review & editing, Funding acquisition; **A. Stier:** Conceptualization, Methodology, Data collection, Data curation, Writing – original draft, Writing – review & editing, Funding acquisition.

## Data availability

All relevant data and details of resources are provided within the article and its supplementary information. Additional materials, including databases and R scripts, are available in vogelwarte.ch Open Repository and Archive (Repository link: LINK PROVIDED UPON ACCEPTANCE).

