## Supplementary Materials for "Slow mitochondria, long life: a phylogenetic comparative analysis in 41 avian species"

**Table S1.** **Details list of the 41 species present in the study.**

| **Scientific name** | **Species** | **Maximal longevity (years)** | **Body mass (g)** | **n** |
| --- | --- | --- | --- | --- |
| *Acanthis flammea* | Common redpoll | 12,2 | 14,2 | 2 |
| *Acrocephalus schoenobaenus* | Sedge warbler | 11,8 | 11,8 | 10 |
| *Acrocephalus scirpaceus* | Common reed warbler | 14 | 12,3 | 3 |
| *Aptenodytes patagonicus* | King penguin | 27 | 13037 | 14 |
| *Chionis minor* | Lesser sheathbill | 24 | 465 | 5 |
| *Chloris chloris* | European greenfinch | 13,6 | 26 | 8 |
| *Coccothraustes coccothraustes* | Hawfinch | 12,6 | 56,6 | 6 |
| *Coloeus monedula* | Jackdaw | 20,3 | 246 | 4 |
| *Coturnix japonica* | Japanese quail | 6 | 115 | 11 |
| *Curruca communis* | Common whitethroat | 8,9 | 17,7 | 8 |
| *Cyanistes caeruleus* | Blue tit | 14,6 | 10,3 | 10 |
| *Dendrocopos major* | Great spotted woodpecker | 12,7 | 75,1 | 8 |
| *Emberiza schoeniclus* | Common reed bunting | 12,2 | 18,4 | 5 |
| *Erithacus rubecula* | European robin | 19,3 | 17,3 | 5 |
| *Ficedula hypoleuca* | European pied flycatcher | 15 | 13,9 | 2 |
| *Fringilla coelebs* | Chaffinch | 29 | 20,7 | 10 |
| *Gallus gallus domesticus* | Chicken | 15 | 780 | 6 |
| *Glaucidium passerinum* | Pygmy owl | 6 | 65,5 | 4 |
| *Hippolais icterina* | Icterine warbler | 10,8 | 13,2 | 2 |
| *Luscinia luscinia* | Trush nightingale | 8,9 | 23,8 | 3 |
| *Macronectes halli* | Northern giant petrel | 35,4 | 4313 | 5 |
| *Muscicapa striata* | Spotted flycatcher | 11,8 | 15,9 | 2 |
| *Parus major* | Great tit | 15,4 | 17,9 | 7 |
| *Passer montanus* | Eurasian tree sparrow | 13,1 | 20,5 | 3 |
| *Periparus ater* | Coal tit | 9,5 | 9,2 | 3 |
| *Phylloscopus trochilus* | Willow warbler | 11,8 | 8,7 | 6 |
| *Procellaria aequinoctialis* | White-chinned petrel | 39 | 1310 | 6 |
| *Prunella modularis* | Dunnock | 20,8 | 20,2 | 2 |
| *Pygoscelis papua* | Gentoo penguin | 41 | 5950 | 6 |
| *Somateria mollissima* | Common eider | 37,8 | 2130 | 8 |
| *Spinus spinus* | Eurasian siskin | 13,5 | 13,2 | 2 |
| *Stercorarius antarcticus* | Brown skua | 28,8 | 1687 | 6 |
| *Strix aluco* | Tawny owl | 22,4 | 520 | 7 |
| *Sturnus vulgaris* | Common starling | 22,9 | 74 | 2 |
| *Sylvia atricapilla* | Eurasian blackcap | 13,8 | 16,7 | 5 |
| *Sylvia borin* | Garden warbler | 24 | 24,6 | 3 |
| *Tachymarptis melba* | Alpine swift | 26 | 102,7 | 6 |
| *Tringa glareola* | Wood sandpiper | 11,6 | 62,5 | 2 |
| *Turdus merula* | Common blackbird | 21,8 | 103,2 | 5 |
| *Turdus philomelos* | Song thrush | 17,7 | 67,8 | 2 |
| *Turdus pilaris* | Fieldfare | 18,1 | 106 | 5 |

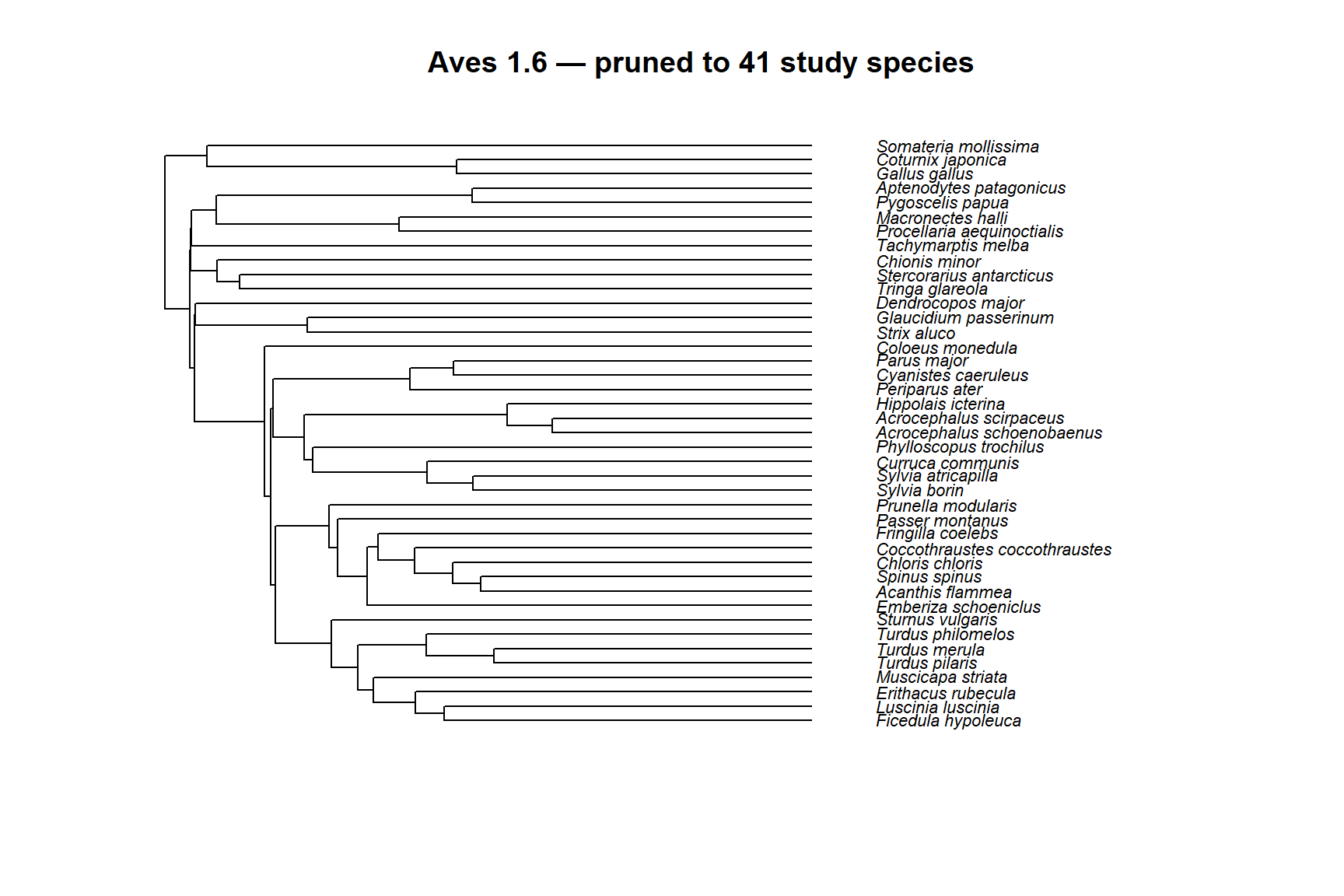

**Figure S1. Phylogenetic tree of the 41 study species (Aves 1.6).** Pruned and ultrametric consensus phylogeny (McTavish et al., 2025) used as the phylogenetic random-effect structure in all MCMCglmm models, showing the evolutionary relationships among the 41 bird species included in this study.

**Table S2.** **Posterior mean estimates and 95% credible intervals (CI) for the effects of log-body mass and log-lifespan on mitochondrial respiration rates and flux control ratios, from phylogenetic mixed models (MCMCglmm).** *P_MCMC_* gives the two-sided Markov chain Monte Carlo p-value for each estimate. Estimates in bold have 95% CI excluding zero. Respiration rates (*ROUTINE*, *OXPHOS_CI_*, *OXPHOS_CI+CII_*, *LEAK*) were estimated jointly in a single multivariate model and are not statistically independent of one another; ratio traits (OxCE, *R/OXPHOS_CI+CII_*, *OXPHOS_CI_* / *OXPHOS_CI+CII_*) were each fitted in separate univariate models. See Methods for model and prior specification. Mitochondrial respirations were scaled prior to modelling.

|  | **log(Body mass)** | | **log(lifespan)** | |
| --- | --- | --- | --- | --- |
|  | Posterior mean [95% CI] | *P_MCMC_* | Posterior mean [95% CI] | *P_MCMC_* |
| **Mitochondrial respiration** |  |  |  |  |
| *ROUTINE* | 0.22 [-0.17; 0.67] | 0.306 | **-3.32 [-4.78; -1.74]** | **<0.001** |
| *OXPHOS_CI_* | **-0.52 [-0.94; -0.08]** | **0.015** | -1.14 [-2.69; 0.40] | 0.147 |
| *OXPHOS_CI+CII_* | **-0.50 [-0.97; -0.08]** | **0.021** | -0.87 [-2.61; 0.73] | 0.292 |
| *LEAK* | **-0.60 [-1.02; -0.20]** | **0.010** | -0.86 [-2.38; 0.68] | 0.274 |
| **Ratios** |  |  |  |  |
| *R/OXPHOS_CI+CII_* | **0.71 [0.32; 1.07]** | **0.001** | **-2.17 [-3.57; -1.03]** | **0.002** |
| *OXPHOS_CI_ / OXPHOS_CI+CII_* | -0.34 [-0.78; 0.13] | 0.133 | -0.76 [-2.43; 0.78] | 0.343 |
| *OxCE* | 0.280 [-0.21; 0.81] | 0.263 | 0.74 [-0.99; 2.54] | 0.407 |

Note: bold indicates estimates whose 95% CI excludes zero. Mitochondrial respiration estimates come from a single multivariate model; ratio estimates come from separate univariate models.

**Table S3.** **Posterior mean estimates and 95% credible intervals (CI) for the effects of log-body mass and residual lifespan on mitochondrial respiration rates and flux control ratios, from phylogenetic mixed models (MCMCglmm).** *P_MCMC_* gives the two-sided Markov chain Monte Carlo p-value for each estimate. Estimates in bold have 95% CI excluding zero. Respiration rates (*ROUTINE*, *OXPHOS_CI_*, *OXPHOS_CI+CII_*, *LEAK*) were estimated jointly in a single multivariate model and are not statistically independent of one another; ratio traits (OxCE, *R/OXPHOS_CI+CII_*, *OXPHOS_CI_* / *OXPHOS_CI+CII_*) were each fitted in separate univariate models. See Methods for model and prior specification. Mitochondrial respirations were scaled prior to modelling.

|  | **log(Body mass)** | | **Residual lifespan** | |
| --- | --- | --- | --- | --- |
|  | Posterior mean [95% CI] | *P_MCMC_* | Posterior mean [95% CI] | *P_MCMC_* |
| **Mitochondrial respiration** |  |  |  |  |
| *ROUTINE* | -0.31 [-0.63; 0.02] | 0.066 | **-3.17 [-4.65; -1.63]** | **<0.001** |
| *OXPHOS_CI_* | **-0.71 [-1.05; -0.38]** | **<0.001** | -1.06 [-2.61; 0.36] | 0.157 |
| *OXPHOS_CI+CII_* | **-0.67 [-1.04; -0.32]** | **<0.001** | -0.64 [-2.14; 1.02] | 0.437 |
| *LEAK* | **-0.73 [-1.06; -0.37]** | **<0.001** | -0.61 [-2.27; 0.87] | 0.438 |
| **Ratios** |  |  |  |  |
| *R/OXPHOS_CI+CII_* | **0.35 [0.05; 0.66]** | 0.030 | **-1.96 [-3.15; -0.72]** | **0.004** |
| *OXPHOS_CI_ / OXPHOS_CI+CII_* | **-0.46 [-0.80; -0.12]** | 0.011 | -0.82 [-2.43; 0.63] | 0.280 |
| *OxCE* | **0.38 [0.005; 0.79]** | **0.051** | 0.75 [-0.97; 2.58] | 0.399 |

Note: bold indicates estimates whose 95% CI excludes zero. Mitochondrial respiration estimates come from a single multivariate model; ratio estimates come from separate univariate models.

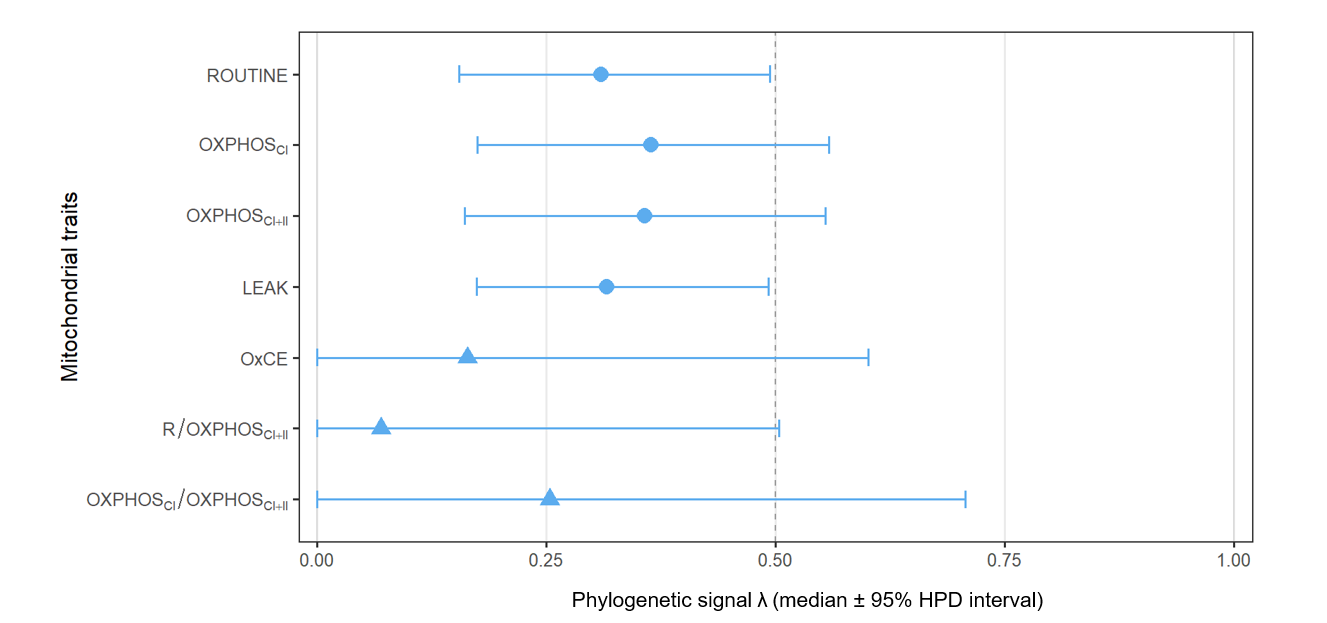

**Figure S2. Phylogenetic signal of mitochondrial traits.** Posterior median phylogenetic signal (λ) and 95% highest posterior density (HPD) intervals for each mitochondrial trait, estimated from multivariate (*ROUTINE, OXPHOS_CI_, OXPHOS_CI+II_,* and *LEAK* shown as circles) and univariate (*OxCE, R/OXPHOS_CI+II_, OXPHOS_CI_/OXPHOS_CI+II_* shown as triangles) MCMCglmm models.

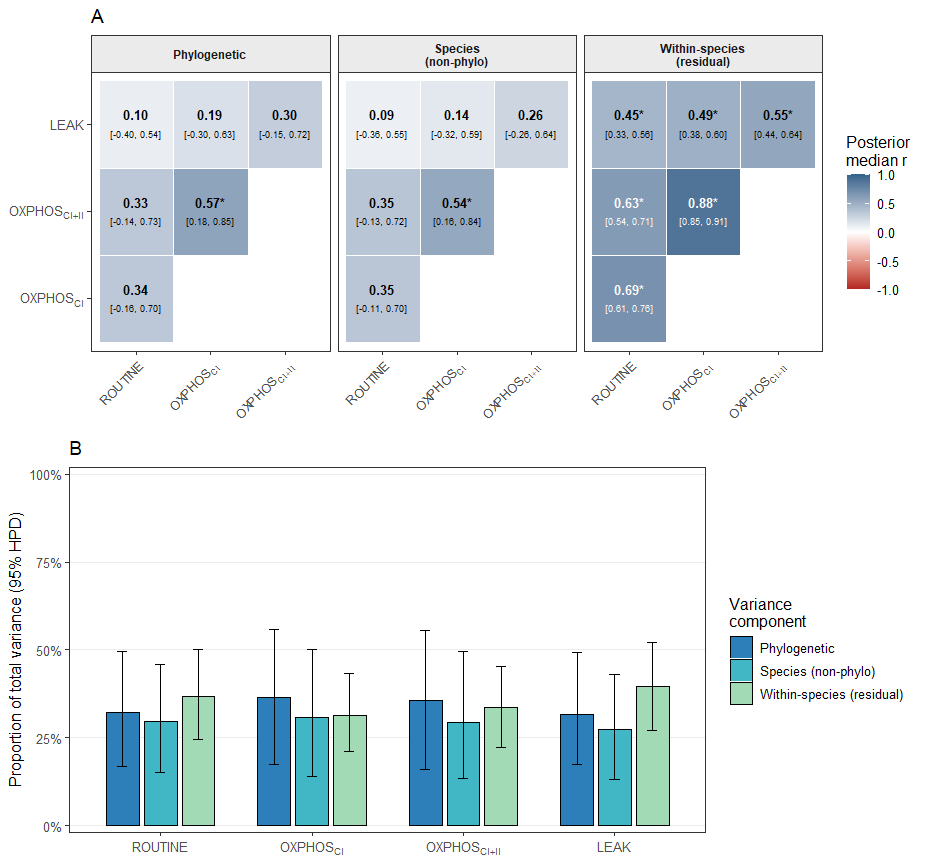

**Figure S3. Phylogenetic, species-level, and residual variance and covariance structure among mitochondrial traits in birds, estimated from a multivariate MCMCglmm animal model. (A)** Posterior median correlation coefficients between *ROUTINE, OXPHOS_CI_, OXPHOS_CI+II_,* and *LEAK*, estimated separately at the phylogenetic level (reflecting evolved covariance among traits attributable to shared ancestry), the species level (reflecting repeatable among-species covariation not explained by phylogenetic relatedness), and the residual/within-species level (reflecting covariation among individuals of the same species). Values show the posterior median r with the 95% highest posterior density (HPD) interval in brackets; asterisks denote pairs whose 95% HPD interval excludes zero. **(B)** Proportion of total among-individual variance attributed to each of the three variance components (phylogenetic, species non-phylogenetic, and within-species/residual) for each mitochondrial trait. Bars show posterior median proportions with 95% HPD intervals; because HPD intervals are computed on the marginal posterior of each ratio independently, bars are not constrained to sum exactly to 100% across components.

*Note: All variance and covariance estimates were obtained from an unstructured (co)variance structure (us(trait)) fitted separately at the phylogenetic, species, and residual levels within a single multivariate MCMCglmm model (family = Gaussian, 1,500,000 iterations, 200,000 burn-in, thinning = 500). Phylogenetic and species-level random effects share the same grouping factor (species identity).*
